# The road not taken: disconnection of a human-unique cortical pathway in schizophrenia and its effects on naturalistic social cognition

**DOI:** 10.1101/2021.08.04.455137

**Authors:** Gaurav H. Patel, David Gruskin, Sophie C. Arkin, Emery C. Jamerson, Daniel Ruiz-Betancourt, Casimir C. Klim, Juan P. Sanchez-Peña, Laura P. Bartel, Jessica K. Lee, Jack Grinband, Antigona Martinez, Rebecca A. Berman, Kevin N. Ochsner, David A. Leopold, Daniel C. Javitt

## Abstract

**Background:** Efficient processing of complex and dynamic social scenes relies on intact connectivity of many underlying cortical areas and networks, but how connectivity deficits affect this functioning in social cognition remains unknown. Here we measure these relationships using functionally based localization of social cognition areas, resting-state functional connectivity, and movie-watching data.

**Methods:** In 42 schizophrenia participants (SzP) and 41 healthy controls (HC), we measured the functional connectivity of areas localized by face-emotion processing, theory-of-mind, and attention tasks. We quantified the weighted shortest path length between visual and medial prefrontal theory-of-mind areas in both populations to assess the impact of functional connectivity deficits on network structure. We then correlated connectivity along the shortest path in each group with movie-evoked activity in a key node of the theory-of-mind network (TPJp).

**Results:** SzP had pronounced connectivity deficits in temporoparietal junction/posterior superior temporal sulcus (TPJ-pSTS) areas involved in face-emotion processing (t(81)=4.4, p=0.00002). In HC the shortest path connecting visual and medial prefrontal theory-of-mind areas passed through TPJ-pSTS, whereas in SzP the shortest path passed through prefrontal cortex (PFC). While movie-evoked TPJp activity correlated with connectivity along the TPJ-pSTS pathway in both groups (r=0.43, p=0.002), it additionally correlated with connectivity along the PFC pathway only in SzP (r_SzP_=0.56, p=0.003).

**Conclusions:** Connectivity along the human-unique TPJ-pSTS pathway affects both the network architecture and functioning of areas involved in processing complex dynamic social scenes. These results demonstrate how focal deficits can have widespread impacts across cortex.

## Introduction

The ability to deftly navigate complex social situations is a key aspect of every day human life (1). During real-life social situations, novel sensory information (e.g. detection of a changing facial expression) is used to update internal models critical to ongoing cognitive operations (e.g. theory-of-mind/mentalization) in order to optimally guide behavior (e.g. who to next look at) (2,3). For the perception and use of these social cues to happen quickly, information must constantly flow within and between the various cortical areas and networks underlying these processes.

Schizophrenia is marked by deficits throughout these processes, including the recognition of facial expressions of emotion, attention, and theory-of-mind operations. Task-based fMRI studies suggest that these deficits are reflected in the abnormal activity across multiple cortical areas, but the interplay between these deficits remains unclear (4,5). Resting-state functional connectivity is often used to assess this degree of functional interaction as it partly reflects how efficiently or frequently any two areas communicate with each other during everyday cognitive processes (6). These relationships can be used to build a “connectome” that maps the cortical network structure (7). The efficiency of communication between any two areas in this map is measured as the shortest path length (7). Therefore, connectivity deficits that affect the shortest path length between two areas may result in abnormalities in the communication between two areas during typical functioning.

A critical step in such analyses is the definition of cortical areas as nodes. Resting-state functional connectivity patterns are often used to parcellate the cortical surface into individual areas, which are combined into networks based on connectivity pattern similarity. These networks are then assigned labels largely based on their resemblance to the spatial patterns of previously described networks derived from task data. While this approach has been revelatory, it also has significant limitations. One is the “reverse inference” problem—the functional interpretation of connectivity deficits is often derived from these connectivity-based network labels, rather than the actual function governed by the areas. This is compounded by the lack of agreement between parcellation schemes, especially in associative regions of cortex. Another limitation is the absence of labels pertaining to specific functions, such as face-emotion recognition, leaving no *a priori* method for identifying and studying these specific areas.

Here instead we use a number of tasks designed to localize the brain areas underlying the key aspects of the perception and use of social cues. These task-based areal definitions provide a more precise functional context within which connectivity deficits can be interpreted as compared to inferring function from network labels(8). We previously used these methods to detail the organization of the temporoparietal junction/posterior superior temporal sulcus (TPJ-pSTS) (8), a region with little consensus about its functional subdivision. In our model, the TPJ-pSTS is the core component of a human-unique pathway linking visual cortex and prefrontal cognitive and theory-of-mind areas that complements the dorsal and ventral visual pathways (2,9). Key components of the TPJ-pSTS include 1) the pSTS, involved in processing moving facial expressions, 2) the TPJp, critical to the performance of theory-of-mind operations, 3) the TPJa, a key node of the ventral attention network involved in re-orienting attention to behaviorally relevant stimuli, and 4) the TPJm, a potential hub linking the other TPJ-pSTS areas.

In a subsequent study (8), we used a naturalistic visual stimulus—a 15-minute clip from the movie “The Good, the Bad, and the Ugly” with the soundtrack removed—to show that in schizophrenia the TPJ-pSTS pathway appeared to be functionally “disconnected”, in that it did not appropriately respond to visual motion inputs, communicate with other visual processing and attention areas, and was not appropriately involved in a key output behavior, saccades. Activation of the TPJ-pSTS was also correlated with performance on a naturalistic test of social cognition, The Awareness of Social Inference Test (TASIT) (10), which we had previously linked to the ability to orient attention/gaze to moving facial expressions in peripheral vision (11).

In this study, we directly examine the connectional integrity of the TPJ-pSTS pathway and its relationship to the naturalistic functioning of a key theory-of-mind area in schizophrenia. We first search for connectivity deficits between these functionally defined areas and component networks. We then measure the impact of these deficits on the structure of the overall network, allowing us to make inferences about how these deficits may impact the functioning of the network. Finally, we test these inferences by examining relationships between resting-state functional connectivity and functioning of the network measured during movie-watching. Based on our previous observations, we hypothesized that a) the structural integrity of the TPJ-pSTS pathway is critical for the intact functioning of key theory-of-mind areas b) that the functional disconnection observed previously is linked to structural deficits along the TPJ-pSTS pathway, and c) that any structural deficits along the TPJ-pSTS pathway would result in shifting the functional reliance of these theory-of-mind areas away from the TPJ-pSTS pathway to alternate processing pathways.

## Materials and Methods

### Participants

42 SzP and 41 HC were recruited with informed consent in accordance with New York State Psychiatric Institute’s Institutional Review Board (IRB). All participants completed the resting-state fMRI portion of the study. The resting-state data reported in (12) represent a subset of these participants. Of these resting-state participants, 27 SzP and 21 HC also completed the movie-watching fMRI session, same as those reported in Patel *et al.* 2020 (8). Inclusion/exclusion criteria are in **Supplemental Materials**.

### Resting-State and Movie-Watching MRI Sessions

Participants were placed comfortably inside the MRI scanner. For the resting-state fMRI scans, participants were instructed to fixate on a centrally placed cross while BOLD data were collected. Multiple 5.5-minue runs of BOLD data were collected (median(interquartile range): HC 4(3.75-5); SzP 4(3-5); t_81_: 0.9, p=.4) using a multiband (MB) fMRI sequence (2mm isotropic, TR=850ms, MB factor 6). Structural T1 and T2 (0.8mm isotropic), along with distortion correction scans (B0 fieldmaps), were also acquired as required for use of the Human Connectome Project (HCP) processing pipelines. Participants willing to return for a second MRI session viewed a video clip of the first 15 minutes of the cinematic movie “The Good, the Bad, and the Ugly” (United Artists, 1966) (13) with the sound removed while BOLD data were collected during one continuous 15-minute acquisition (1049 MR frames) with the same parameters as the resting-state fMRI scans.

### Image Processing

MRI data were preprocessed with the HCP pipelines v3.4 (14) which place the data into gray-ordinates in a standardized surface atlas (as opposed to voxels in a volume atlas). The functional data were additionally cleaned of artifact largely following the recommendations from Power *et al.* (15). An adaptive filter was used to censor movement-related frames to minimize the effects of respiratory motion (16). The percentage of frames censored differed by a small but significant amount (mean(sd); HC: 14.6% (5.4%) versus SzP 20.1% (8.1%); t_81_=3.6, p=0.0006).

### Localizing Regions of Interest

TPJ-pSTS ROIs were derived from three localizer tasks: 1) a task that contrasted activity evoked by moving versus static facial expressions, designed to localize areas involved in face-emotion recognition (17); 2) a visual search task designed to activate areas involved in visual processing, visual attention, and cognitive control (12,18,19), and 3) a task designed to activate areas involved in theory-of-mind operations evoked during the viewing of a short animated video (20). Based on the localizers and prior studies, these 98 ROIs were then subdivided into 28 component systems. Details about these tasks, how they were used to define the ROIs, and how they were divided into component systems are in **Supplemental Materials** and have been described previously in Patel *et al.* 2021 (8) for the localization of TPJ-pSTS ROIs.

### Contrasting Resting-State Functional Connectivity Between SzP and HC

Pairwise resting-state functional connectivity was calculated by correlating the cleaned time-course of each ROI pair (Pearson correlations) within each participant. Between and within-component connectivity was calculated by averaging ROI-ROI connectivity pairs within each component-component interaction. Matrices for each individual were then Fisher-z transformed and used to calculate both within-group averages and between group contrasts using two-tailed t-tests (Bonferroni-corrected for multiple comparisons). Custom scripts implemented in MATLAB (Natick, MA) were used for this and all subsequent analyses.

### Network Structure and Efficiency Analyses

To quantify network structural and efficiency differences between groups, thresholded group-average component connectivity matrices were used to calculate the weighted shortest path length between L early visual (EVis) cortex and the R antero-medial theory-of-mind areas (amToM, involved in long timescale theory-of-mind predictions (3)) after inversion of the connectivity weights to represent inter-nodal distance (Brain Connectivity Toolbox, https://sites.google.com/site/bctnet/). This revealed two potential paths, one running through the TPJ-pSTS and the other through prefrontal cortex (PFC).

To calculate group differences in the paths connecting L EVis and R amToM areas, we first repeated the above shortest weighted path length calculation for each individual. We then created two “gates” from the component nodes, one representing the TPJ-pSTS pathway (L and R pSTS) and the other representing the PFC pathway (L and R PFC and L and R cingulo-opercular/salience or COSal). We then calculated the percentage of individuals whose L EVis to R amToM paths included the TPJ-pSTS gate nodes and the PFC gate nodes. This calculation was repeated after thresholding the connectivity matrices for every percentile threshold between the 50^th^%tile and the 85^th^%tile (above which the number of individual graphs with no path between visual cortex and amToM areas began to increase precipitously) and repeating the shortest path length and gate analyses. No restriction was placed on the shortest path length search to force inclusion of one or the other gate. We then calculated the 2×2 contrast of the percentage of paths through the TPJ-pSTS versus PFC gates for SzP versus HC for all thresholds by comparing the area under the curve (AUC) for each of the four measures. We used permutation testing to calculate the significance of these comparisons by shuffling the group labels and recalculating each measure 10,000 times, creating a null distribution against which to compare the actual AUC differences. All analyses were repeated for R EVis to R amToM paths with similar results.

### Correlation of Connectivity with Path Length

To examine the relationship of specific ROI-ROI connectivity strengths and path lengths, we used Pearson correlation after a Fisher-z transformation of the connectivity matrices. For the specific L EVis to R amToM paths tested above, we co-varied for group to determine whether the correlation was similar or different in the two groups.

### Correlation of Connectivity to TPJp Movie-Evoked Activity

For the subset of individuals who performed the movie-watching experiment, we first calculated the amount of movie-evoked activity in TPJp using inter-subject correlation (ISC) (13). TPJp ISC values were calculated for each participant as the pairwise correlation of the TPJp ROI time-course for each individual and the TPJp time-courses of each HC participant. HC ISC values were calculated against the group of all other HCs. We then correlated ROI-ROI connectivity strengths with TPJp ISC values, co-varying for group.

## Results

### Demographics

Demographically, SzP were matched to HC, with a small increase in mean(sd) age (39.8(10.8) vs. 34.9(9.6), p=.03) in SzP (see **Table 1**). As shown previously in a sub-sample of these participants (11), SzP were more impaired in TASIT Sarcasm versus TASIT Lie performance and on the Processing Speed Index. Other demographics and SzP characteristics are in **Table 1**. For the subset of participants who performed the movie-watching experiment, no significant demographics differences were found (8).

**Table 1:**
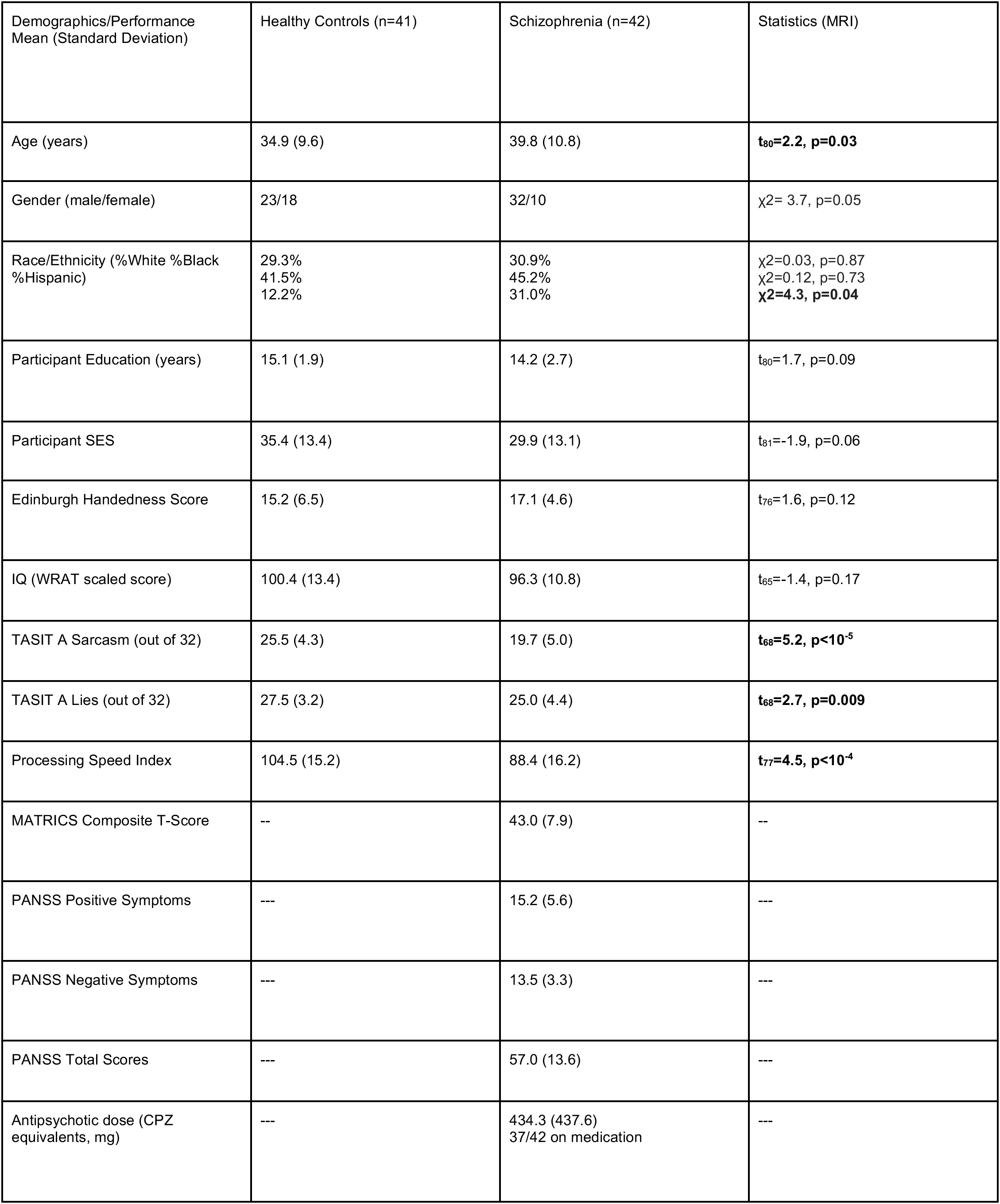
Demographics, scales, and test performance.

### Parcellation of Brain Areas Activated by TASIT and the Movie-watching Task

Localizer tasks were used to subdivide the cortex activated by TASIT into functionally defined parcels, grouped together into component networks that included visual processing, dorsal/ventral attention, face/face-emotion recognition, prefrontal, cingulo-opercular/salience, and mentalization/theory-of-mind component networks (colored borders in **Figure 1A** and **Supplemental Figure 1**). A similar pattern of activity (as measured by ISC) was evoked by watching the longer visual-only movie clip (**Figure 1B)**, with (as expected) the most significant discrepancy in the auditory cortex and superior temporal gyrus (STG).

**Figure 1:**
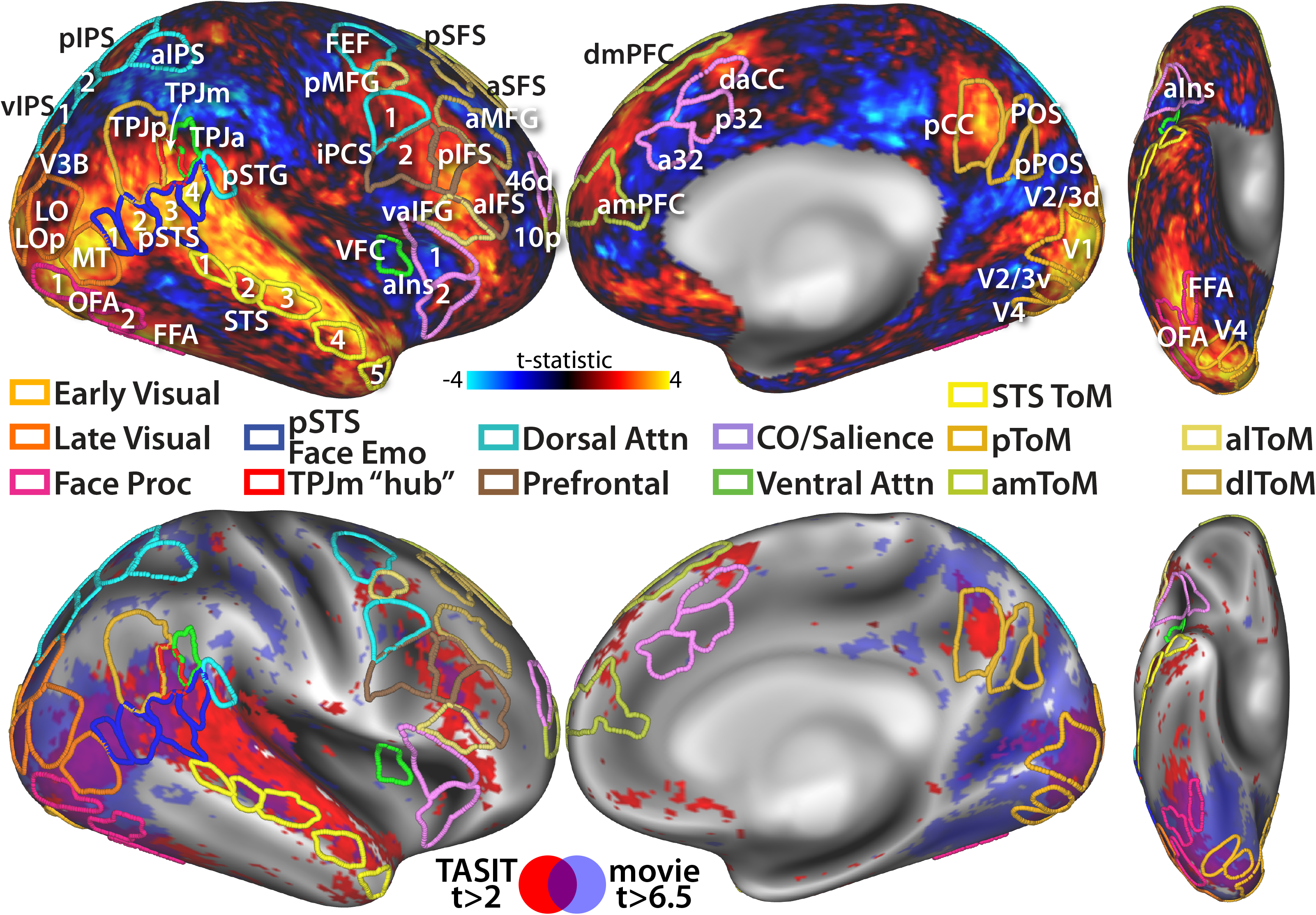
Cortical activation by naturalistic stimuli and region of interest (ROI) definitions. **A:** Activation evoked by TASIT videos in 10 HC are covered by ROIs derived from functional localizers (with the notable exception of auditory cortex). See **Supplemental Figure 1** for functional localizers and ROI definitions. **B:** Activation by TASIT (red) and activation by the cinematic movie “The Good, the Bad, and the Ugly” as measured by intersubject correlation (ISC, blue) largely overlap (purple). *V2/3v:* ventral V2 and V3 *LO:* lateral occipital *LOp:* lateral occipital posterior *MT:* medial temporal *OFA:* occipital face area *FFA:* fusiform face area *pSTS:* posterior superior temporal sulcus *STS:* superior temporal sulcus *pSTG:* posterior superior temporal gyrus *TPJa:* temporoparietal junction anterior *TPJm:* temporoparietal junction middle *TPJp:* temporoparietal junction posterior *vIPS:* ventral intraparietal sulcus *pIPS:* posterior intraparietal sulcus *aIPS:* anterior intraparietal sulcus *FEF:* frontal eye-fields *pMFG:* posterior middle frontal gyrus *iPCS:* inferior precentral sulcus *VFC:* ventral frontal cortex *aIns:* anterior insula *p/aSFS:* posterior/anterior superior frontal sulcus *aMFG:* anterior middle frontal gyrus *aIFS:* anterior inferior frontal sulcus *vaIFG:* ventral anterior inferior frontal gyrus *10p:* posterior area 10 *46d:* dorsal area 46 *dmPFC:* dorsal medial prefrontal cortex *amPFC:* anterior medial prefrontal cortex *a32:* area 32 *daCC:* dorsal anterior cingulate cortex *pCC:* posterior cingulate cortex *POS:* parietal/occipital sulcus *pPOS:* posterior POS *Proc:* Processing *Emo:* Emotion *Attn:* Attention *CO:* cingulo-opercular *ToM:* theory-of-mind *pToM:* posterior ToM *amToM:* anteromedial ToM *alToM:* anterolateral ToM *dlToM:* dorsolateral ToM

### Functional Connectivity of TASIT-activated Areas

Across all of the component systems, we observed significant deficits in SzP in the within-system connectivity of the pSTS face-emotion recognition and lateral/ventral occipitotemporal face-processing systems (see **Figure 2A** and **Supplemental Figures 2-3**). Focusing on these two components revealed an additional strong difference between the late visual and pSTS components (**Figure 2B**). **Supplemental Figures 4-5** demonstrates the same general pattern of deficits with the parcellation scheme from Glasser *et al.* (21). No significant correlations of age or other demographics with connectivity in the pSTS or other areas was observed.

**Figure 2:**
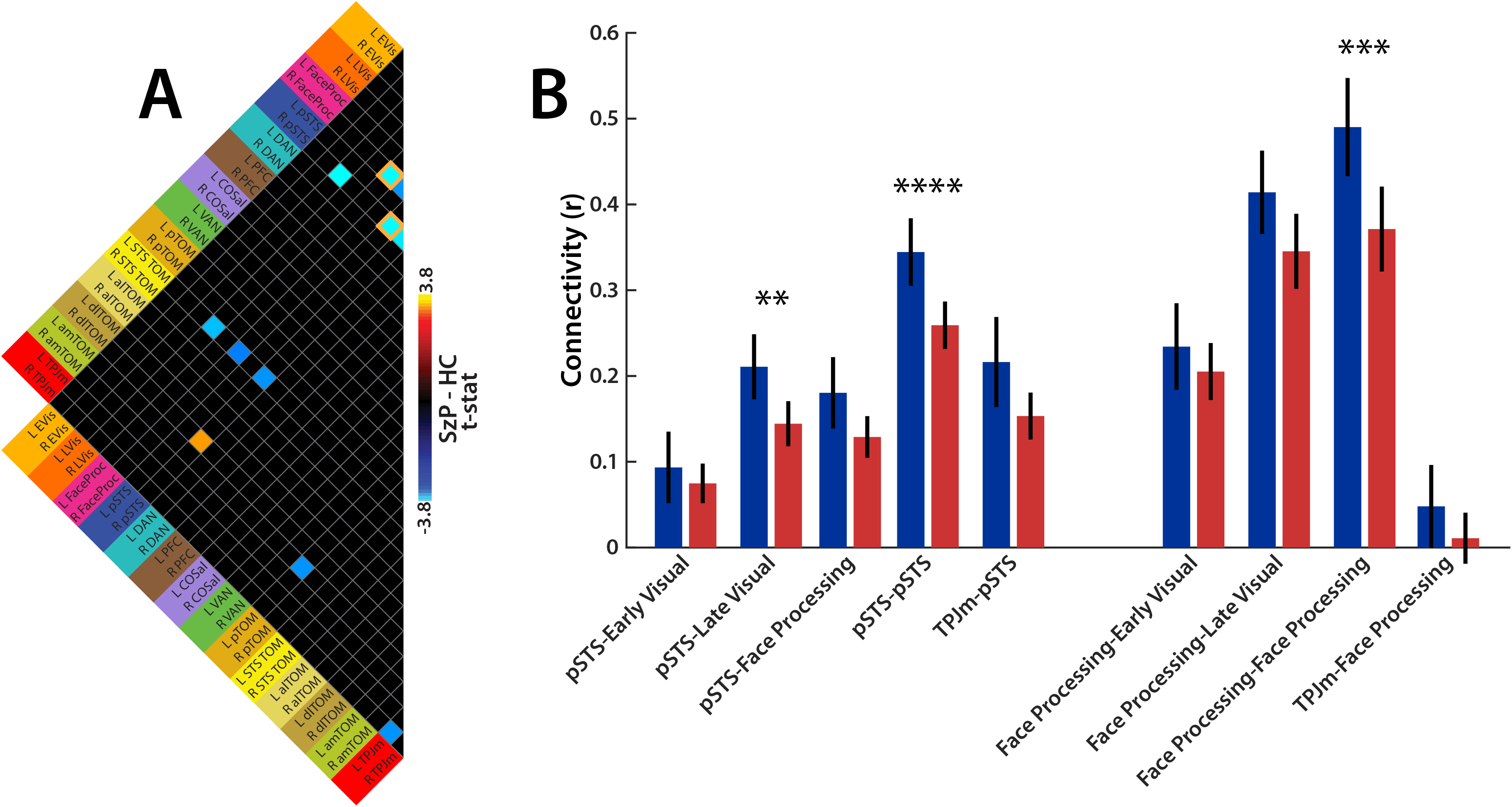
Contrast of component-to-component connectivity between SzP and HC. **A:** Differences between SzP and HC component connectivity are limited to the face-processing and pSTS components. Matrix thresholded at p<0.01. Orange outlines highlight differences that survive multiple comparisons at p<0.05: face-processing: t(81)=4.3, Cohen’s d=0.95; pSTS: t(81)=4.4, Cohen’s d=0.96). **B:** Detail of differences in SzP versus HC connectivity of the face-processing and pSTS components with themselves, visual, and TPJm components. All starred differences significant after correction for the 9 comparisons made. **p<0.01 ***p<0.001 ****p<0.0001

### Reduced TPJ-pSTS Pathway Efficiency and Increased PFC Pathway Efficiency in SzP

We next examined the effects of the connectivity deficits on network structure and efficiency. In **Figure 3A**, visual areas are closely connected to dorsal attention areas which in turn are closely connected to prefrontal and cingulo-opercular areas. Some prefrontal/cingulo-opercular areas in turn are connected to the theory-of-mind areas, forming one potential path connecting visual areas to theory-of-mind areas (PFC pathway, highlighted in orange). The TPJ-pSTS areas appear to form a separate path connecting visual to theory-of-mind areas (TPJ-pSTS pathway, highlighted in green), containing the affected connections detailed above.

**Figure 3:**
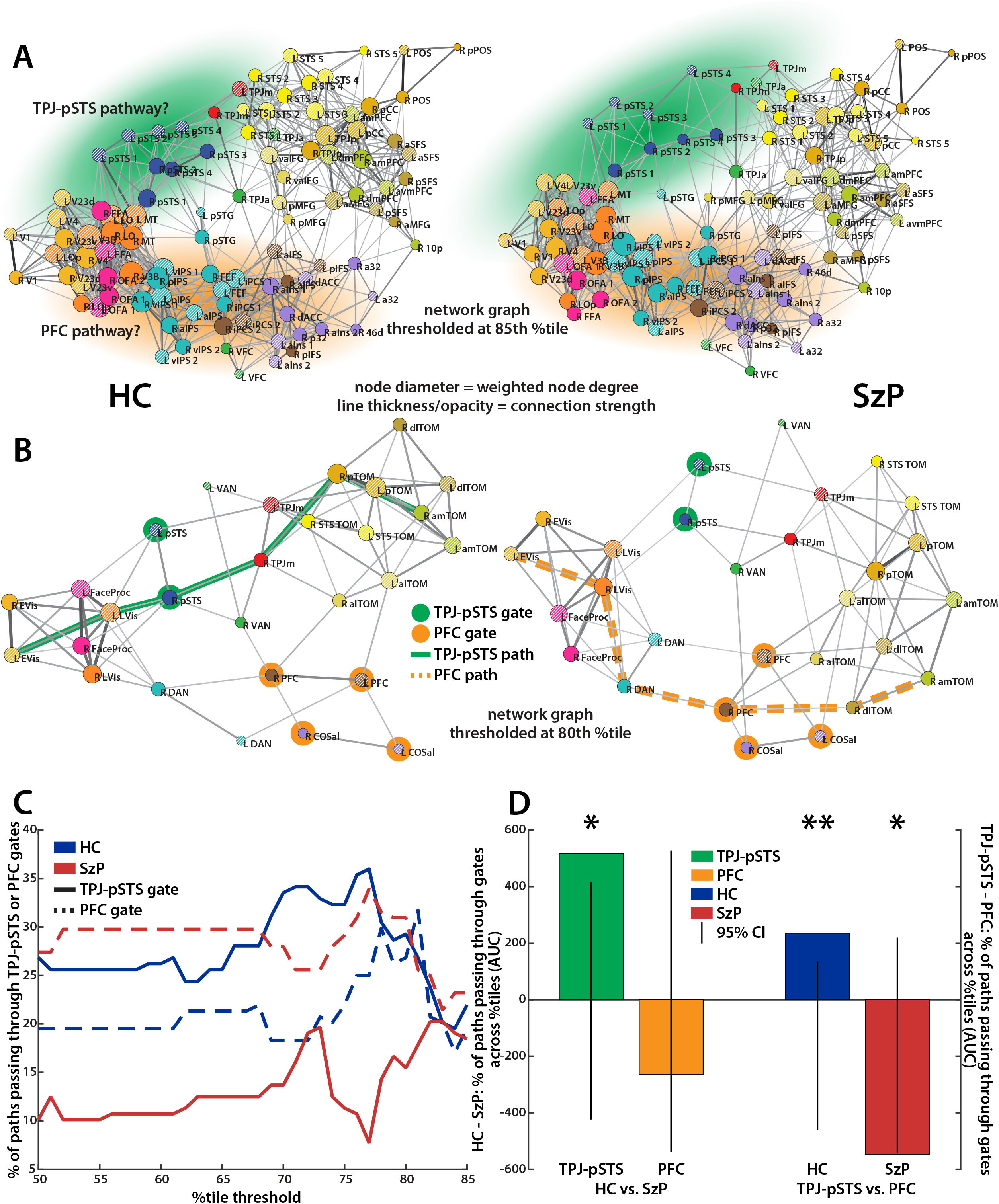
Structural changes caused by pSTS deficits as measured by the path length between early visual (EVis) cortex and right medial prefrontal cortex (amToM). **A:** ROI-ROI network graphs for HC and SzP. Spring-loaded graphs were produced in Pajek (http://mrvar.fdv.uni-lj.si/pajek/) and used to visualize the network architecture ROIs. The nodes in the graphs represent ROIs and are arranged such that the distance between any two nodes in the spring-loaded graph is inversely proportional to the similarity in their connectivity patterns with other areas. The connectivity strength between nodes is represented by the thickness of the line connecting the nodes, and the number and strength of connections to that node (weighted node degree) is represented by the diameter of the nodes. Weighted node degree was chosen as it provides information about which components are most critical for the functioning of a network (41,42). ROI-ROI group-average connectivity matrices were thresholded at the 85^th^%tile before network graph visualization, and the Kamada-Kawai free-energy algorithm was used to arrange the ROIs. Component color codes are the same as **Figure 1**. Green shading highlights potential TPJ-pSTS pathway and orange shading the potential PFC pathway. **B:** Component-component network graphs for HC and SzP in roughly the same spatial configuration as **Figure 3A** but with nodes representing components instead of individual ROIs and group-average connectivity matrices thresholded at the 80^th^%tile. Green line highlights the shortest weighted path between L EVis and R amToM components in HC. This path runs through the TPJ-pSTS “gate” (green highlighted nodes) and is hence labeled the TPJ-pSTS pathway. Orange dotted line highlights the shortest weighted path between L Evis and R amToM components in SzP. This path runs through the the PFC “gate” (orange highlighted nodes) and is hence labeled the PFC pathway. **C:** The percentage of paths in individual participants connecting L/R EVis and R amToM in HC and SzP that run through the TPJ-pSTS gate nodes (highlighted in green) and/or the PFC gate nodes (highlighted in orange) as a function of the percentile threshold applied to create the graph. **D:** The difference in the area under the curve (AUC) for each of the four comparisons represented in **Figure 3C** compared to a null distribution (represented by the line). Between groups, significantly more paths across HC participants run through the TPJ-pSTS than in SzP (green bar) whereas the percentage of paths running through the PFC is similar in the two groups (orange bar). Within HC, significantly more paths run through the TPJ-pSTS than the PFC (blue bar). Within SzP, significantly more paths run through the PFC than TPJ-pSTS (red bar). *p<0.05 **p<0.01

To examine how the pSTS deficits in SzP affected the shortest path between visual and theory-of-mind areas, we first compared the shortest weighted path length between the L Evis and R amToM component systems (**Figure 3B**). In HCs this path ran through the TPJ-pSTS pathway and included the late visual cortex, TPJm, and the posterior subdivision of the theory-of-mind network (which includes the area TPJp on the angular gyrus). In SzP, this path instead ran through prefrontal cortex (orange dotted line) and included components of the PFC pathway (dorsal attention, prefrontal, and the dorsal lateral prefrontal subdivision of the theory-of-mind areas (dlToM). This group difference was similar across percentile thresholds (**Figure 3C**), with the shortest path crossing significantly more often through the TPJ-pSTS in HCs vs. SzP (**Figure 3D** green bar). Within groups, the shortest path in HCs was most often along the TPJ-pSTS pathway (**Figure 3D** blue bar) versus the PFC pathway in SzP (**Figure 3D** red bar).

### Group Differences in the TPJ-pSTS and PFC Integrity and Pathway Efficiency between Visual and Theory-of-Mind Components

We then examined whether the connectional integrity along the TPJ-pSTS and PFC pathways affected the path length between visual and theory-of-mind areas in one or both groups. Here, a negative correlation means that *stronger* component-component functional connectivity is correlated with *shorter* path length between L Evis and R amToM, which in turn would suggest that the connectivity between those components was important for the communication efficiency between visual and theory-of-mind areas. Along the TPJ-pSTS pathway, connectivity of L LVis to R pSTS correlated strongly with L Evis and R amToM path length across groups (r=−0.46, **Figure 4A**). This relationship was particularly pronounced in SzP versus HC (r_HC_=−0.37 versus r_SzP_=−0.59). This association generalized to the path length between any visual/face processing/face-emotion component and any theory-of-mind component along the TPJ-pSTS pathway (**Figure 4B** left side, green labels) but not to PFC components (orange labels). A similar pattern was observed for R LVis to R pSTS connectivity (**Figure 4B** right side).

**Figure 4:**
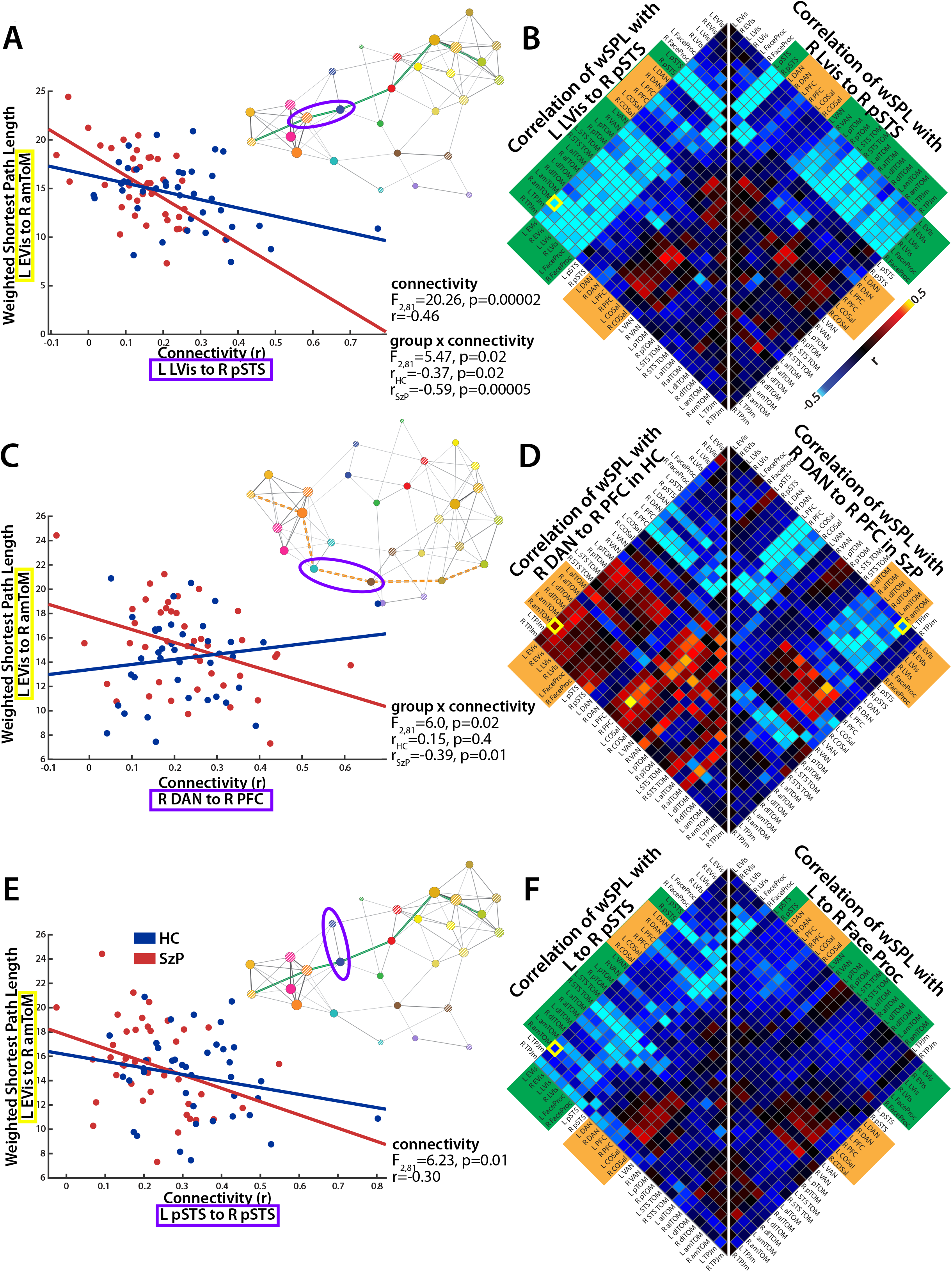
Group differential effects of TPJ-pSTS and PFC pathway integrity on component-component path efficiency. **A:** Connectivity strength between L LVis and R pSTS components (purple ellipse on network graph) affects L EVis to R amToM weighted shortest path length (wSPL) in both groups. Negative correlation indicates that stronger connectivity leads to shorter (more efficient) path lengths. **B:** Connectivity strength between L LVis and R pSTS components (left half) affects component-component path length of TPJ-pSTS pathway components (green path on network graph, matrix labels highlighted in green) and not with path lengths between components along the PFC path (labels highlighted in orange). R LVis and R pSTS component-component connectivity has a similar relationship with component-component path length (right half). Correlation in **Figure 4A** highlighted in yellow. **C:** Connectivity strength between R DAN and R PFC components (purple ellipse on network graph) along the PFC pathway (orange dotted line on network graph) affects L EVis to R amToM path length in SzP but not HC. **D:** Connectivity strength between R DAN and R PFC components affects component-component path length between visual and theory-of-mind areas in SzP (right half, labels highlighted in orange) but not HC (left half). Correlation in **Figure 4C** highlighted in yellow. See **Supplemental Figures 6-7** for other TPJ-pSTS and PFC path components and at other percentile thresholds. **E:** Connectivity strength between L pSTS and R pSTS components (purple ellipse on network graph) affects L EVis to R amToM path length in both groups. **F:** The relationship of connectivity strength between L pSTS and R pSTS components and component-component path length (left half) affects primarily TPJ-pSTS pathway components (green line on network graph, green matrix labels) over PFC pathway components (orange labels) and is similar to L/R LVis to R pSTS connectivity in **Figure 4B**. The relationship of connectivity strength between L face-processing and R face-processing components and component-component path length, however, only affects path length between face-processing and visual areas (right half). Correlation in **Figure 4E** highlighted in yellow.

Along the PFC pathway, connectivity of the R DAN to R PFC correlated with decreased path length between L EVis to R amToM in SzP but not in HC (r_HC_=0.15 versus r_SzP_=-0.39, **Figure 4C**). This dissociation between groups generalized to the connectivity between visual/face-processing areas and prefrontal theory-of-mind-areas (**Figure 4D**, orange labels), with HC having no or weakly positive relationships (**Figure 4D** left side) and SzP having strongly negative relationships (**Figure 4D** right side). See **Supplemental Figures 6 & 7** for connectivity × path length correlation matrices for other components of the TPJ-pSTS/PFC pathways at various percentile thresholds.

### Effects of the Functional Connectivity Deficits on Pathway Efficiency between Visual and Theory-of-Mind Components

We then examined whether the pSTS and/or face-processing component connectivity deficits affected the integrity of one or both pathways. R to L pSTS connectivity was correlated with L EVis to R amToM path length across groups(r=−0.30, **Figure 4E**). This relationship generalized to the path length between any visual/face processing/face-emotion component and any theory-of-mind component (**Figure 4F**, left half). No similar relationship existed for connectivity between R to L face-processing areas (**Figure 4F**, right half). The pSTS deficit correlation pattern was significantly more similar to the L LVis to R pSTS pattern in **Figure 4B** than the face-processing deficit correlation pattern was to the L LVis to R pSTS pattern (r=0.83 versus r=0.51, p<10^−16^), suggesting that only the pSTS deficits affected the intergirty of the TPJ-pSTS pathway.

### Relation of Pathway Connectivity to the Functioning of Theory-of-Mind Areas during Naturalistic Stimulation

The above connectivity-path length analyses suggest that pSTS deficits in SzP affect functioning of the TPJ-pSTS pathway and increase use of the PFC pathway. In the subset of participants with movie-watching data, we tested this hypothesis by examining relationships between the connectivity of component nodes along each pathway with the functioning of a key theory-of-mind area, the TPJp (**Figure 5**). The connectivity deficits detailed in **Figure 2** replicate in this subsample (**Supplemental Figure 8**). Along the TPJ-pSTS pathway, connectivity of early/late visual and face-processing areas to the TPJm (where SzP had the largest functional deficits in Patel *et al.* 2021 (8)) positively correlated in both groups with activation of the TPJp (**Figure 5A**, r=0.43). Conversely, connectivity of R pSTS to R amToM negatively correlated with TPJp activation in both groups (**Figure 5B**, r=−0.53), with the strongest activation of TPJp associated with negative connectivity. While a similar overall pattern is observed with the R TPJm to R amToM connectivity (**Figure 5C**, r=−0.50), there is a difference between groups with a significant negative correlation in HC (r=−0.74) but not SzP (r=−0.15).

**Figure 5:**
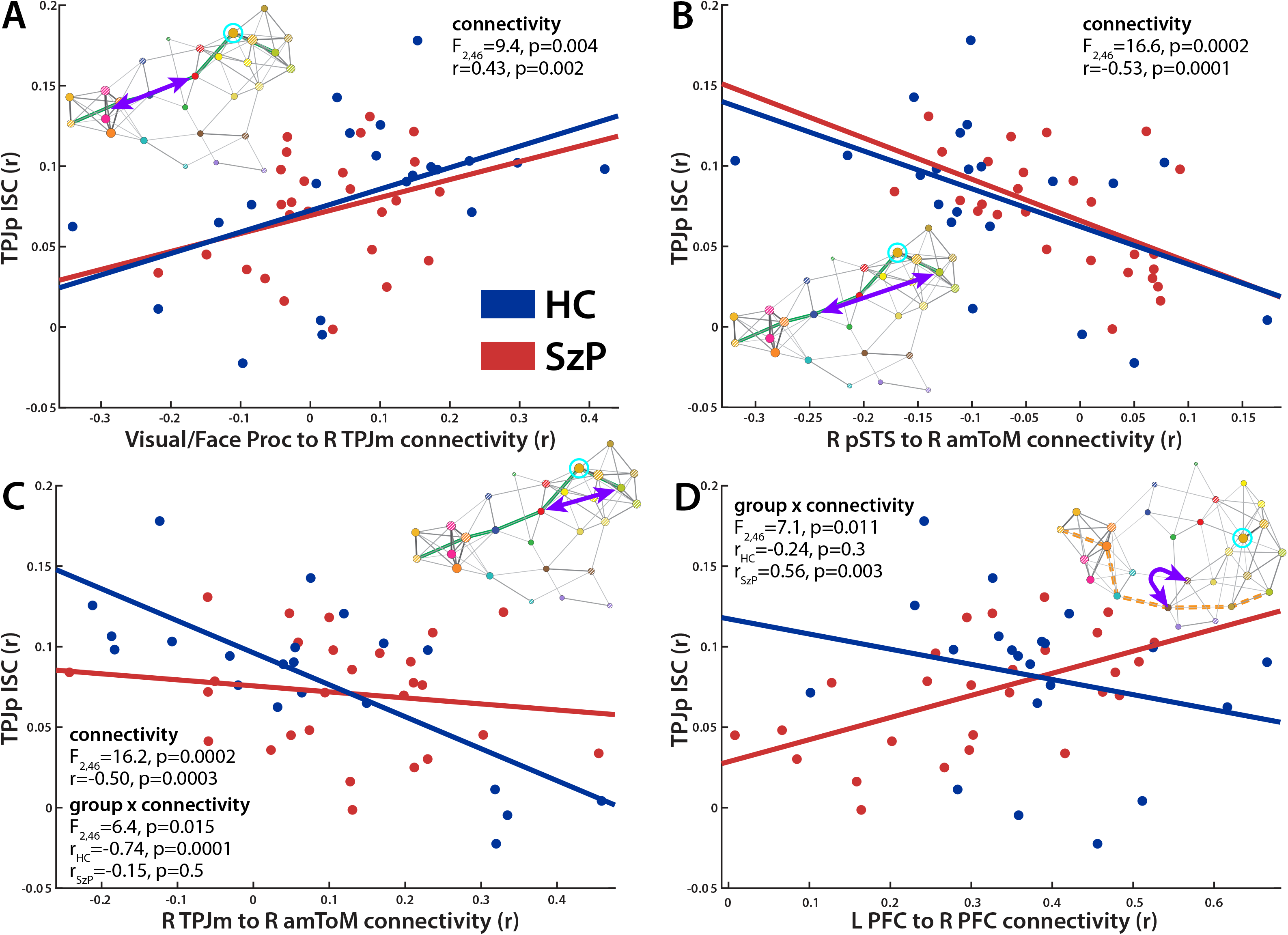
Relationship between connectivity along the TPJ-pSTS and PFC paths and TPJp activation as measured by ISC. **A:** Connectivity of visual/face-processing areas to TPJm positively correlates with TPJp activation similarly across groups. **B:** Connectivity of R amToM to pSTS negatively correlates with TPJp activation similarly across groups. **C:** Connectivity of R amToM to TPJm negatively correlates with TPJp activation in HC but not SzP. **D:** Connectivity between L and R PFC positively correlates with TPJp activation in SzP but not HC.

Along the PFC pathway, we observed that connectivity between L and R PFC predicted TPJp activity in SzP but not HC (**Figure 5D**, r_HC_=−0.24 versus r_SzP_=−0.56, p=0.003). See **Supplemental Figure 9** for connectivity and group × connectivity F statistics for other TPJ-pSTS and PFC pathway components.

## Discussion

In this study, we examined the functional connectivity of areas involved in the processing of naturalistic social scenes in SzP versus HC. First, we found focal deficits in SzP involving the pSTS face-emotion processing areas. Second, we found that the most efficient path connecting visual areas to medial prefrontal theory-of-mind areas involved two separable pathways in the two groups—the TPJ-pSTS pathway in HC and the PFC pathway in SzP. Third, we found that in HC efficient connectivity between visual and theory-of-mind components relied on the integrity of the TPJ-pSTS pathway, whereas in SzP this connectivity relied on the integrity of both the TPJ-pSTS and PFC pathways. Fourth, we found that the pSTS deficits primarily affected connectivity of visual and theory-of-mind areas along the TPJ-pSTS pathway. These connectivity analyses suggested that the pSTS deficits in SzP shifted the functional reliance of key theory-of-mind areas away from communication along the TPJ-pSTS pathway and towards the PFC pathway. We then confirmed these hypotheses by finding that the fidelity of the movie-evoked activation of the TPJp, a key theory-of-mind area, depended on the TPJ-pSTS pathway integrity in both groups, but on the PFC pathway integrity only in SzP. Overall, these results demonstrate that not only can focal functional connectivity deficits result in network-wide structural changes with downstream functional impacts, but also point to potential mechanisms of resilience in the face of a major deficit.

### Comparison to Previous Studies

Previous functional connectivity studies of SzP have not found the deficits revealed here (22–25), but our study is unique in two key ways. First, ours is one of the first to use high-resolution multiband sequences and surface mapping methods developed for the Human Connectome Project. This is of particular importance for the TPJ-pSTS, where high-degrees of individual variability in sulcal geometry may obscure or blur potential differences in group averages and comparisons (26–28). Second, previously published parcellation schemes not only disagree greatly in the parcellation of the TPJ-pSTS, they also do not subdivide the TPJ-pSTS by function, with no designated face-emotion processing and attention reorienting areas (2). The resulting connectivity deficits complement previous studies of visual processing and face emotion recognition in schizophrenia. SzP have long been noted to have a range of visual processing deficits (29–33) including in face-emotion recognition (34,35) and the processing of biological motion (36). These deficits have been linked to the lateral occipital cortex for visual processing (31,32) and the pSTS for face-emotion recognition (17). The functional connectivity deficits in this study likely reflect those deficits, and may represent disorganization or decreased/impaired use of this cortical region (37,38). Of note, the connectivity deficits described in (12) overlap with those discussed here.

### TPJ-pSTS as a Separate Processing Pathway

The observation that variation in TPJ-pSTS pathway connectivity only affects the path lengths between other components of the TPJ-pSTS pathway (but not PFC pathway components) supports our theory that the TPJ-pSTS represents a separate processing pathway linking visual and theory-of-mind areas, distinct from the classical dorsal/ventral visual processing pathways (both embedded in the PFC pathway) (2). As detailed in that review, this human-unique pathway allows for internal models of mental states to be more quickly updated by relevant incoming sensory information about social cues than the dorsal/vental visual processing pathways embedded in the PFC pathway. This fast updating then leads to more efficient planning of eye-movements and other relevant social behaviors. The movie-watching analyses confirm that efficient transmission of the visual information through the pSTS into the TPJ is critical for “normal” activation of the TPJp, a key area involved in maintaining and updating these internal mental state models.

### Effects of pSTS Deficits on Connectivity-Function Relationships in SzP

The pSTS connectivity deficits observed here in the SzP effectively lesion the TPJ-pSTS pathway, seemingly preventing its use in processing the naturalistic stimuli. These results suggest that the TPJ deficits observed previously by our group (8) were downstream of the pSTS face-emotion processing deficit as opposed to being rooted in the TPJ itself. The failure of prefrontal-TPJm connectivity to predict TPJp ISC further supports the idea that the TPJm has been removed from the processing pathway in SzP.

These functional deficits (reflected by connectivity deficits) along the TPJ-pSTS pathway also appear to shift the reliance of TPJp activation away from just the TPJ-pSTS pathway. Our analyses suggest that in SzP the PFC pathway, which contains the classically defined dorsal/visual streams, serves as an alternate processing pathway for conveying visual information to theory-of-mind areas, with increasing connectivity along this pathway predicating improved fidelity of TPJp activation. The idea of there being multiple pathways for these operations has to be true of course: macaques are still able to visually scan social scenes and perform limited theory-of-mind operations despite having only the PFC pathway but not the TPJ-pSTS pathway (39).

The fact that PFC pathway connectivity predicts TPJp activation *despite the TPJp not being a member of this pathway* is particularly interesting, and may complement a number of other observations we have made. First, in Patel *et al.* 2021 (8), we observed that in SzP the TPJ was less synchronized with visual and dorsal attention areas but more synchronized with prefrontal theory-of-mind and cognitive control areas than in HC, suggesting that the disconnection from visual processing areas had resulted in *increased* communication with prefrontal areas. Second, we observed that in SzP (but not HC) only the Processing Speed Index (PSI) from the WAIS-III IQ test strongly predicted TASIT performance (11). The tasks that compose the PSI all involve some degree of top-down driven visual search which is guided by dorsolateral prefrontal cortex, the same areas that correlate with TPJp performance in this manuscript. These observations suggest that in SzP the more visual information enters the theory-of-mind network through prefrontal cortex, the more “normal” the activity is in the theory-of-mind areas such as TPJp. However, the visual strategy enacted by use of the PFC pathway could also result in differences in the relationship between visual scanning and theory-of-mind operations, as we observed recently (11).

### Limitations and Future Directions

These results complement those reported in previous studies from our group (8,11) from overlapping samples and demonstrate several previously unreported organizational principles and relationships; therefore, the results will need to be replicated in an independent sample. Larger samples will also allow for the study of how intervening levels of analysis (task activation, low-level behaviors) mediate the relationship between resting-state functional connectivity and high-level behaviors linked to symptoms. Studies like this may also provide insight into the small effect sizes observed when relating resting-state functional connectivity and behavioral measures (40). Another aspect of the TPJ-pSTS that merits further investigation is the extent to which individual variation in areal boundaries affects the structural-functional relationships discussed here. Perhaps most importantly, the model of the interactions within the TPJ-pSTS and across networks proposed here will need to be tested with causal methods, such as transcranial magnetic stimulation (TMS). Nonetheless, the results shown here and in our related studies provide a framework of how areas within the TPJ-pSTS and across cortex dynamically update models of other people’s mental states in social situations with incoming sensory information to guide future behaviors. With this framework, we will be better able to guide both the creation of both diagnostic biomarkers of social dysfunction and the development of treatment targets aimed at alleviating the suffering from these deficits.

## Acknowledgements

We wish to thank the funding agencies who supported this work: NIMH (GHP: K23MH108711, T32MH018870, R01MH123639, and R01MH121790; DCJ: R01MH049334; DAL and RAB: Intramural Research Program ZIA MH002898), Brain & Behavior Research Foundation (GHP), American Psychiatric Foundation (GHP), Sidney R. Baer Jr. Foundation (GHP), Leon Levy Foundation (GHP), the Dana Foundation (GHP), and the Herb and Isabel Stusser Foundation (DCJ).

## Data Availability

Data will be made available after reasonable request to corresponding author.

