## Supplemental Material for "The road not taken: disconnection of a human-unique cortical pathway in schizophrenia and its effects on naturalistic social cognition"

### Supplementary Methods

*Inclusion/Exclusion Criteria for Participants*

Participants were primary English speakers, ages 18-55 with no history of head trauma or neurological disorders and with normal or corrected visual acuity (20/40 or better). Participants with contraindication for MRI (artificial implants and/or claustrophobia) were excluded from the study. Patients met DSM-IV criteria for schizophrenia, as confirmed by the Lieber Schizophrenia Research Clinic (LSRC) before recruitment and participation. Patients were on disease-appropriate medication used within appropriate guidelines, and were assessed with the Positive and Negative Symptoms Scale (PANSS; (1)). Patients scoring above 120 on the PANSS, or those who were acutely psychotic were excluded. HC were recruited through the LSRC and through IRB-approved flyers and internet advertisements. The Structured Clinical Interview for the DSM-IV (SCID-NP) was used to exclude past or present Axis I or II disorders, significant substance use disorders in the past 6 months, and in HC significant psychiatric family history (2).

*Apparatus and Procedural Setup for fMRI*

For resting state, task, and movie-watching experiments, participants were placed on the scanner bed with cushioning to stabilize potential head movement and earplugs to block out MR-related sound. A mirror attached to the head coil allowed participants to view visual stimuli displayed on a translucent screen mounted at the end of the bore from a liquid crystal display (LCD) projector; the eye-to-screen distance (mirror-to-screen + eye-to-mirror) was 107 cm. Participants held a fiber optic, five-channel button box (Current Designs, Philadelphia, PA) in their right hands that allowed them to respond to target stimuli with their index finger if the task required it. The experimental display was controlled by a MacBook Pro (Apple Inc., Cupertino, CA) running custom software within either MATLAB (Mathworks, Natick, MA) using Psychophysics toolbox version 3.012 (3,4) or Experiment Builder (SR Research, Mississauga, Ontario, Canada), which connected to the projector via high-definition multimedia interface (HDMI) hub. The stimuli were synchronized to the beginning of the MR volume acquisition via a USB device that triggered the MATLAB paradigm to begin (Current Designs, Philadelphia, PA). Eye movements were recorded using Eyelink 1000 plus (SR Research, Mississauga, Ontario, Canada).

*fMRI Acquisition*

Functional and anatomical data were acquired with a 32-channel phased array receive-only head coil (Nova Medical, Wilmington, MA) by General Electric's Discovery MR750 3.0 Tesla full body MR scanner (GE; Fairfield, CT) at New York Psychiatric Institutes' (NYSPI) MRI Research Unit. Localizer scans and a gradient echo B0 fieldmap (TE=4.8ms, TR=800ms, FOV=256mmx256mm, slice thickness=2mm, matrix=128x128, slices=68) were required for the Human Connectome Project (HCP) processing pipelines. Structural images were acquired in 5-minute sequences and were comprised of two T1-weighted images (3D sagittal, 0.8mm isotropic, matrix size=300x300, slices=220, TR=7856ms, TE=3108ms, flip angle=12, TI=450ms) and two T2-weighted images (3D sagittal, 0.8 mm isotropic, matrix size=320x320, slices=220, TR=2500ms, TE=95.708, flip angle=90°). Task and resting state functional data were collected with a multiband SMS-EPI sequence (2 mm isotopic, slice plane=transverse, TR=850ms, MUX=6, ARC=1, TE=25ms, matrix size=96x96, slices=66, phase encoding direction=P→A) (courtesy of the Center for Cognitive and Neurobiological Imaging, Stanford University, http://cni.stanford.edu).

*Resting State Scans*

Participants were asked to fixate on a central point for the duration of the 5.5 minute scan. Runs in which the participant fell asleep (assessed by watching their eyes on the eye-tracker monitor) were excluded from analyses.

*Attention Localizer*

Attention areas were localized with an RSVP visual search task similar to the one used by Patel *et al.* (5,6). 34 HCs performed this task. Forty-two clipart images of neutral objects were used as both target and distractor stimuli. One of the forty-two clipart images was randomly selected as the target image, while the other forty-one images were used as distractors. Before the BOLD run started, the target image was presented once in each of the three stream locations to familiarize the participant with the target. One stream was superimposed over the fixation point in the center of the screen with a width/height of 3.6º. The two peripheral streams were located at 5.8º eccentricity with a polar angle of 0º (left of the fixation point) and 180º (right of the fixation point) and with a width/height of 5.3º. Targets were chosen randomly, under the condition that no target could be used twice in a row. Participants pressed a specified button on a button box with their right index finger every time they saw the target stimulus.

Each trial consisted of a 10 second RSVP stream in one of three locations and with the same sizes described above. Each stimulus in the RSVP stream was displayed for 100 milliseconds (ms) before being replaced by the next distractor with no gap between presentations. Distractor stimuli were chosen randomly with the constraint of not having consecutive repeats. The target stimulus took the place of a distractor stimulus in the stream and did not differ in any parameters from the distracting stimuli. Half of all trials had a single target presented randomly within the first 8 seconds in the 10-second stream. One-quarter of trials had no target (catch trials). In the remaining one-quarter of trials, the target appeared twice within the target stream, once within the first 4 seconds and again from 4 to 8 seconds. After the target appeared, participants had 1.3 seconds to indicate detection. Participants were instructed to respond as accurately and quickly as possible, with an emphasis on accuracy so as to minimize false positives. Following each 10-second stream, the participant was asked to maintain fixation for inter-trial intervals of 2, 4, or 6 seconds, chosen randomly, before the next trial began. Two BOLD-fMRI runs were collected, each 5 minutes 30 seconds each (~20 trials / run). These are the same task data and results reported in (7).

*Static/Moving Emotional Faces Localizer*
To localize areas involved in the processing of static and moving facial expressions, participants passively viewed blocks of static or moving emotional or neutral stimuli (8). The four block types were: 1) “neutral static”, five pictures of the faces of differing people displaying no emotion; 2) “neutral moving”, five videos of the faces of differing people moving but displaying no emotion; 3) “emotion static”, five pictures of the faces of differing people displaying a certain emotion (happiness, anger, fear, sadness); 4) “emotion moving”: five videos of the faces of differing people displaying the same emotions. All stimuli were in grayscale. Pictures were displayed for 2 seconds each, and videos for 4 seconds each, with no gap between either. Blocks were repeated in this order, and interspersed with 20s of fixation. Two runs of 7 minutes 5 seconds each were collected from 24 HC.

*Motion Localizer*
To identify the motion-sensitive visual areas, we used a previously described motion localizer (9,10). In short, participants maintained fixation on a central cross while a series of low-contrast concentric rings of a diameter of 15 degrees expanded or contracted for 20 seconds. These blocks alternated with 20 second blocks during which the same rings remained static. Participants were instructed to respond to the occasional dimming of the fixation point via a button press to maintain attention and fixation. One run of 5 minutes 23 seconds each were collected from 14 HC.

*Theory-of-Mind (Mentalization) Localizer*
To identify areas involved in theory-of-mind operations, we used a previously described movie-watching task (11). Participants were instructed to watch the short, animated clip of Disney Pixar’s “Partly Cloudy”, and then briefly summarize the plot afterwards to ensure attention. A single run of 6 minutes 36 seconds was collected for 12 HCs.

*Movie-watching Task*
Participants viewed the first 15 minutes of the cinematic movie “The Good, the Bad, and the Ugly” (United Artists, 1966) with the audio track silenced while eye-movements were recorded. Participants were asked to briefly summarize the clip afterwards to ensure attention. BOLD-fMRI data were collected in a single run of 1068 frames.

*Image Processing*
All imaging data were processed on workstation machines (Mac Pro, Apple Inc., Cupertino, CA) using the HCP processing pipeline v3.4 adapted for the NYSPI GE MR750 MRI scanner (12). The HCP pipelines first processed the anatomy images to create a cortical surface model for each individual aligned to the HCP fs_LR 32k atlas, then for the functional runs it performed movement correction, distortion correction, and atlas alignment in a single resampling step, and lastly projected the functional data to an atlas cortical surface through the individual’s cortical surface model. After each step, manual quality checks were performed to

check for errors in atlas-alignment and segmentation. This pipeline is different from other processing pipelines in that it creates a Connectivity Informatics Technology Initiative (CIFTI) file for each BOLD run that only contains the data from the cortical and the subcortical grey matter ("gray-ordinates" as opposed to voxels), which allows for a more precise localization of brain activity that is not confounded by cerebrospinal fluid (CSF) or white matter partial volume effects. Structural and functional images were aligned in a volume space of the Montreal Neurological Institute (MNI152) atlas and on the surface Conte69/fs_LR 32k atlas created by the HCP pipeline developers (12,13).

Additional post-processing procedures, adapted from Power et al. (14), were performed to further minimize noise in the resting state and movie-watching data. First, estimates of head motion calculated in the X, Y, and Z directions along with the displacements of rotation around the X, Y, and Z axes (pitch, yaw, and roll) by the HCP movement correction algorithm, along with their derivatives (backwards difference), the square of the six parameters, and the sum of these six parameters (assuming a 50mm radius of head size for the rotations, previously labeled as the framewise displacement (FD)) were used as nuisance regressors to remove movement-related artifact. Second, tissue signals and their derivatives calculated by averaging signal across voxels within a spatial mask for ventricle signals, CSF signal, white matter signal, and whole brain signal were used as nuisance regressors to remove physiology-related noise (global signal regression also removes movement-related artifact (9). Third, MR frames with an FD greater than the 75%tile + 1/2 * interquartile range of the FD trace for each run, limited to a range between 0.2mm and 0.5mm were censored, and replaced by interpolation using a method based on the Lomb-Scargle periodogram (14,15). Fourth, data were low band-pass filtered at .588 Hz (the Nyquist limit for the MR sampling rate) and high band-pass filtered at 0.0005 Hz (12). The resulting time-series was used for all subsequent analyses. All analyses were repeated with the FD threshold set to 0.2mm and with no whole brain signal regression.

*Task fMRI Analyses*
The task fMRI analyses for the attention, motion, and static/moving faces paradigms were all conducted using FEAT (16). For level 1 analyses, the magnitudes of activation evoked by attention to/processing of paradigm-related "events" were separately estimated in each individual via a general linear model (GLM) by modeling each event type as an independent regressor (17). For each paradigm, the regressor was formed by convolving a boxcar function corresponding to the event length with an estimate of the hemodynamic response to an impulse function (18). A high-pass filter was of the length of the BOLD run was used to remove linear trend. The first four volumes of each run were excluded from the analyses. Results from individuals with two BOLD runs were combined in a level 2 fixed-effects analysis. Level 3 group analyses were performed using FLAME 1 mixed-effects.

For the attention localizer, the magnitudes of activation evoked by attention to and processing of the RSVP stream and detection of the target at each spatial location were separately estimated in each individual. For the RSVP stream, the event duration was 10 seconds. For detection events, the reaction time of the response to each target was used as the event duration. For the misses and false detections, the event duration used was 500ms. For the static/moving faces localizer, the modeled event blocks were neutral moving, neutral static, emotion static, and emotion moving. The event durations used for convolution were 20 seconds for the moving and 10 seconds for the static. For the motion localizer, the only modeled event was for the blocks of the expanding/contracting rings, which had a length of 12 seconds.

To analyze the theory-of-mind localizer, we utilized the hand-coded mentalization, social, physical pain and control regressors for the animated short as described in Jacoby *et al.* (11). The four regressors were then convolved with the same estimate of the hemodynamic response to an impulse function used above (Grinband *et al.*, 2017). A one-sample t-test was performed in MATLAB (Natick, MA) to create the statistical map of activations.

*ROI Definitions*
In general, regions of interest (ROIs) were defined on the average activation maps from the task localizers on the cortical surface by drawing borders around each activation based on the activation gradient maps (similar to the procedures using in the creation of the HCP atlas (19). Activation gradient maps derived from the spatial derivative of the z-statistic maps were used to define borders around local maxima (peaks), and to separate contiguous activations by tracing the line that represented the highest gradient (local minimum) between the

two activations. ROI labels were largely based on sulcal/gyral anatomy, with the HCP's multi-modal parcellation of the cortical surface serving as a guide for labeling some of the ROIs.

**Supplementary Figure Captions**

**Supplemental Figure 1:** Localizer cortical task activation patterns and borders. **A)** Motion localization: gray-ordinates activated by motion (moving concentric rings). **B)** Theory of mind (ToM) localization: gray-ordinates correlated with higher-order ToM operations. Yellow ToM borders based these activation maps and then subdivided based on anatomical location (posterior, superior temporal sulcus (STS), anterolateral, anteromedial, and dorsolateral). **C)** Static emotional faces localization: gray-ordinates activated by passive viewing of pictures of facial expression of emotion. Ventral occipito-temporal face-processing borders (pink) based on the resulting activation patterns. **D)** Visual processing and attention localization: gray-ordinates activated during the processing of and attention to the rapid serial visual presentation (RSVP) stream of pictures of objects while searching for the target object. Borders for early visual (light orange), late visual (dark orange), dorsal attention (cyan), prefrontal (brown), and cingulo-opercular/salience (light purple) based on positive activations and separated into components based on matches in location to resting-state networks and task literature. Early/late visual area borders confirmed with motion localizer (**A**). **E)** Moving emotional faces localization: gray-ordinate-wise contrast of activity evoked by the passive viewing of short movies of moving facial expressions versus static emotional faces in **C**. Blue posterior STS borders based on this activation pattern. **F)** Target detection activation: gray-ordinates activated by the detection of the target stimulus in the RSVP task. Green ventral attention borders based on the conjunction of gray-ordinates de-activated during RSVP search (**D**) and activated by target detection (**F**), limited to the posterior aspect of the supramarginal gyrus for the TPJa and ventral frontal/insular cortex for the VFC.

**Supplemental Figure 2:** Component connectivity matrices for HC (left) and SzP (right), along with the group contrast (bottom right).

**Supplemental Figure 3:** Node connectivity matrices for HC (left), SzP (right), along with the group contrast (bottom right).

**Supplemental Figure 4:** TASIT activation pattern parcellated by the Human Connectome Project parcellation scheme (Glasser *et al.* 2016) compared to the task-localized borders used in this study.

**Supplemental Figure 5:** Connectivity matrices for HC (left), SzP (right), and contrast (bottom right) for all Glasser parcels activated by TASIT (**Supplemental Figure 4**). Connectivity deficits largely limited to visual and posterior STS parcels; parcels overlapping with the blue pSTS borders from the moving emotional face localizer highlighted in gray.

**Supplemental Figure 6:** Correlation of connectivity strength of TPJ-pSTS pathway components and the path length between L Evis and R amToM components calculated after thresholding the adjacency matrix at three different percentile cutoffs. Negative correlations reflect stronger connectivity strengths corresponding to shorter path lengths. Patterns for each connectivity-path length relationship are similar across percentile thresholds, including the effect of LVis-pSTS connectivity strength being limited to only TPJ-pSTS pathway components (second row).

**Supplemental Figure 7:** Correlation of connectivity strength of PFC pathway components and the path length between L Evis and R amToM components calculated after thresholding the adjacency matrix at three different percentile cutoffs. Negative correlations reflect stronger connectivity strengths corresponding to shorter path lengths. Patterns for each connectivity-path length relationship are similar across percentile thresholds, including the differences between groups in the relationship in R DAN-R PFC connectivity and visual-ToM component path length (third row).

**Supplemental Figure 8:** Comparison of the group contrast in component connectivity for the full sample (left) and the movie-watching sub-sample (right). The two matrices are very similar (r=0.82).

**Supplemental Figure 9:** Relationship of component connectivity with TPJp ISC after co-varying for group identity. **A** shows TPJ-pSTS pathway components and **B** the PFC pathway components. Left half shows the main effect of the connectivity-ISC relationship, or where the relationship is similar across groups. The right half the group x connectivity interaction, or where the relationship differs between groups. In the TPJ-pSTS pathway, visual to TPJm connectivity and pSTS/TPJm to ToM components show significant relationships across groups, positive correlations for visual to TPJm and negative for pSTS/TPJm to ToM.  However, R TPJm to amToM connectivity also shows a significant difference between groups.  In the PFC pathway, visual to DAN connectivity is significantly related to TPJp ISC in both groups, but there are group differences in the L PFC to L/R PFC connectivity and TPJp ISC relationship.

Supplemental Figure 1

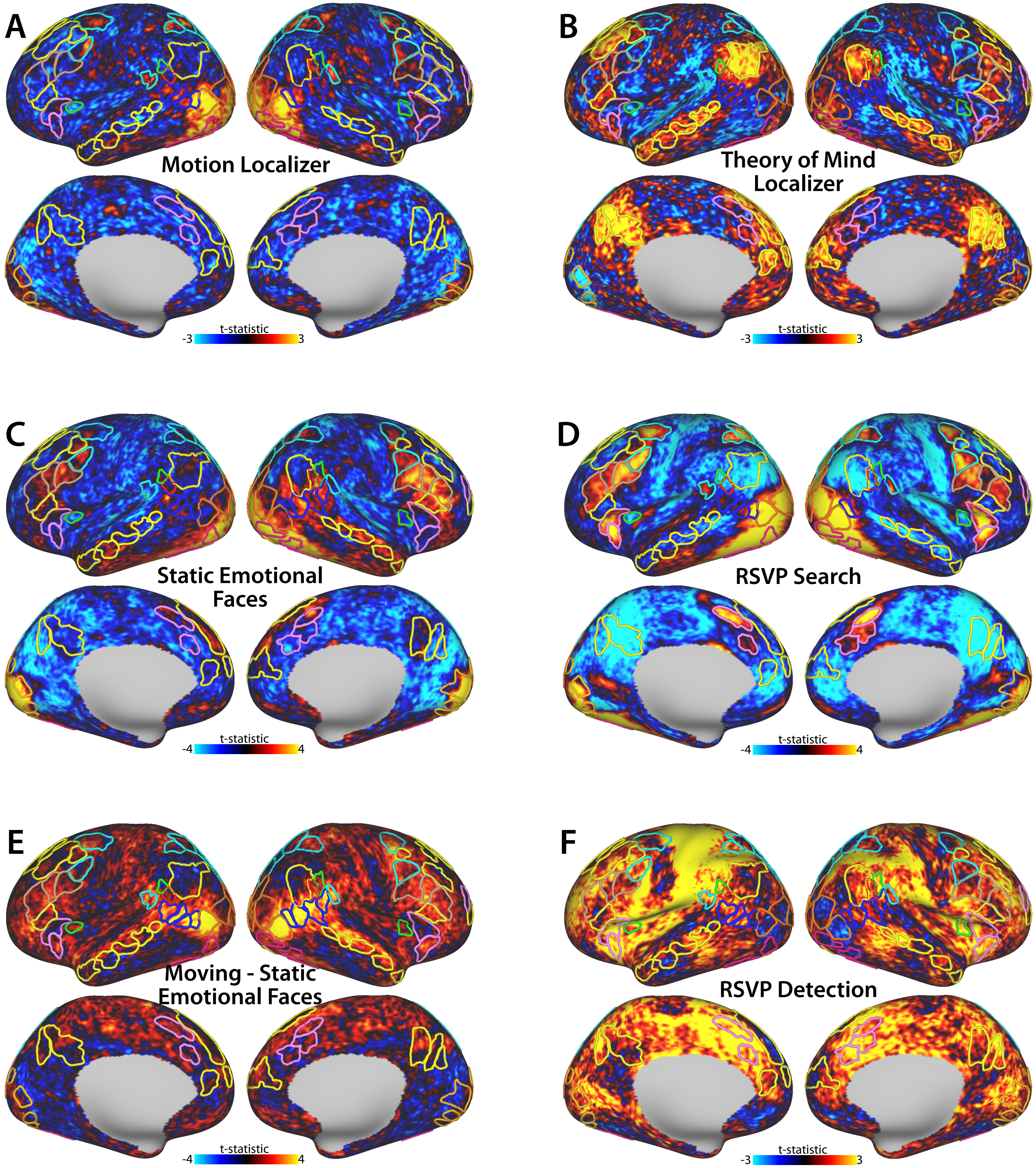

**Supplemental Figure 2**

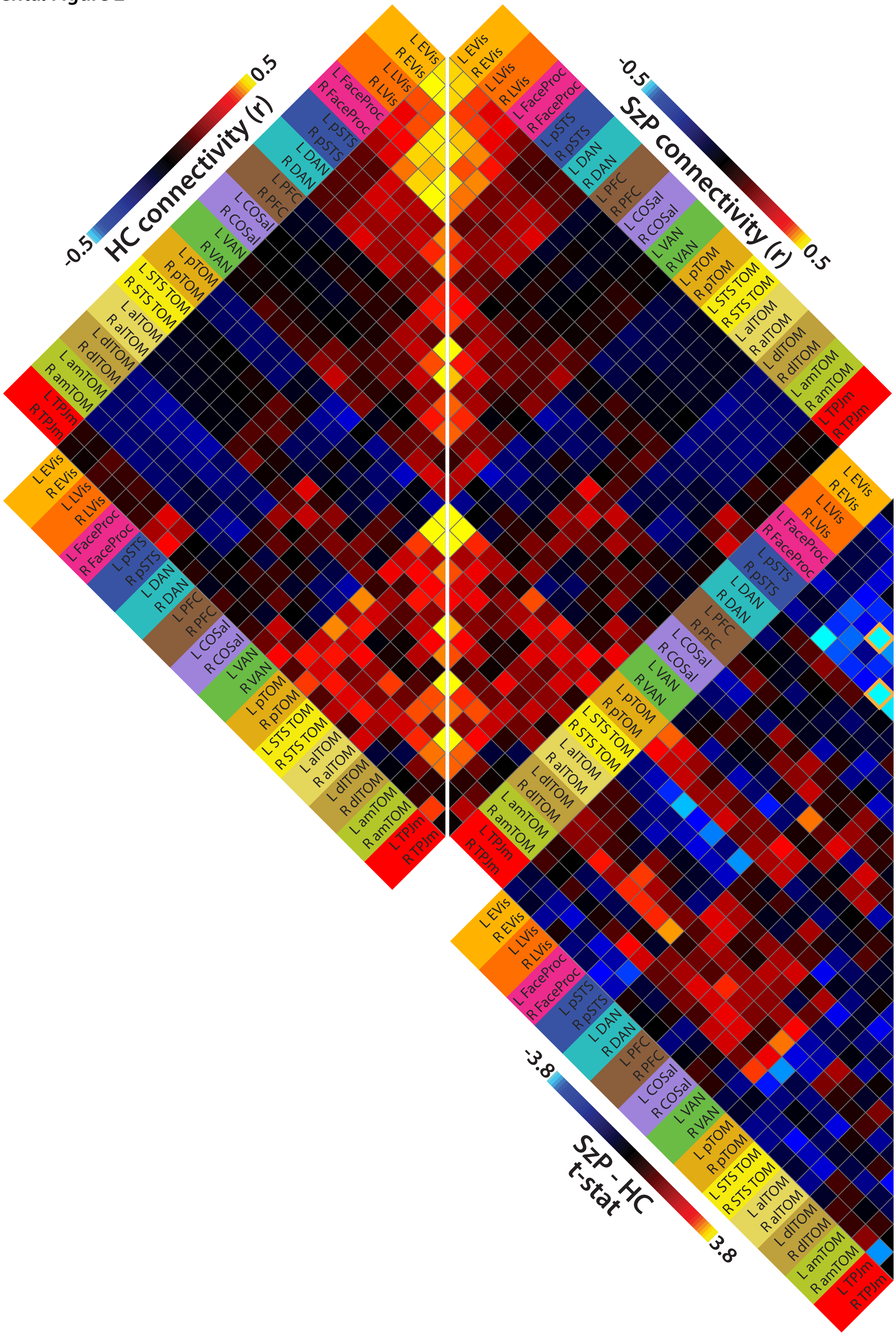

Supplemental Figure 3

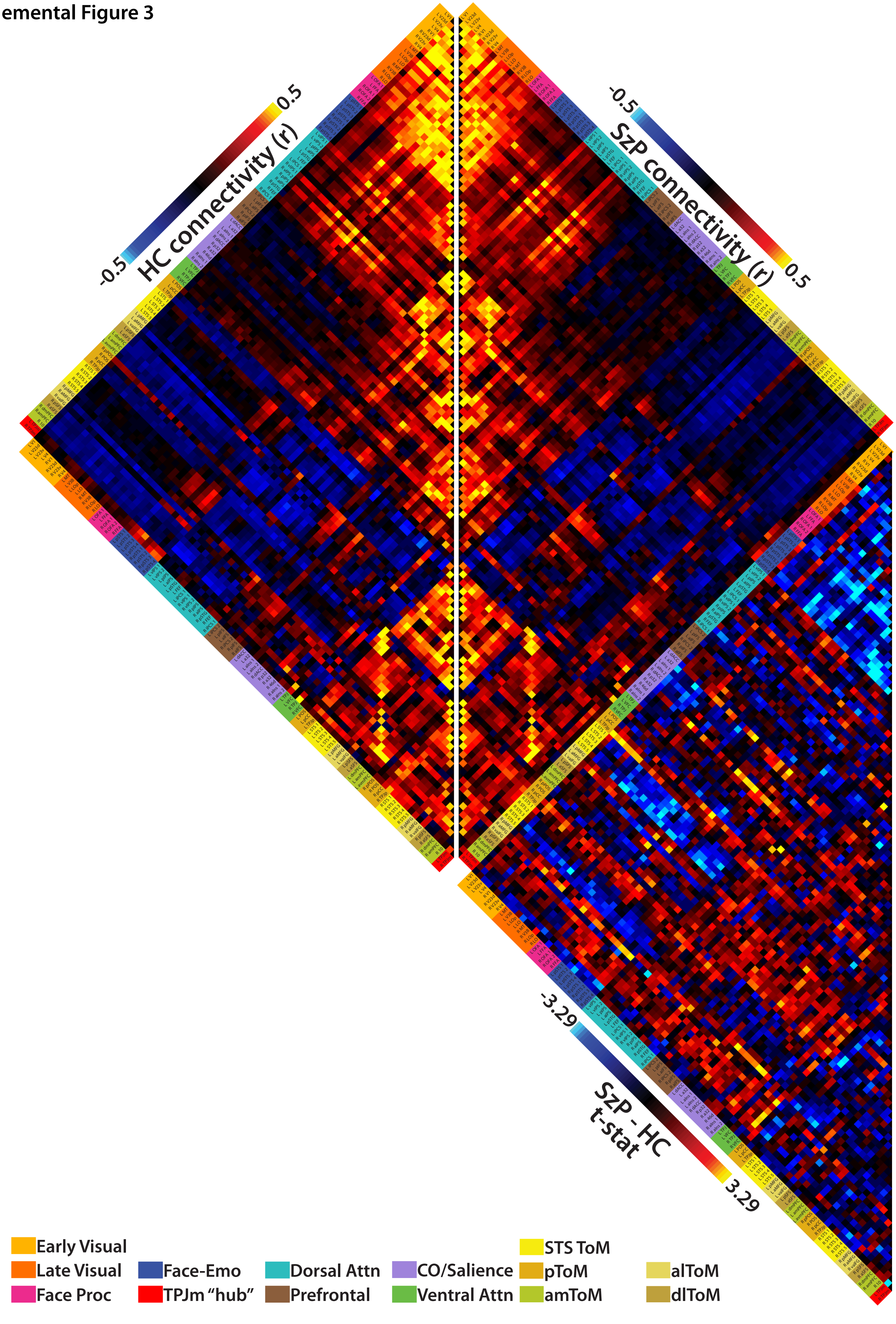

**Supplemental Figure 4**

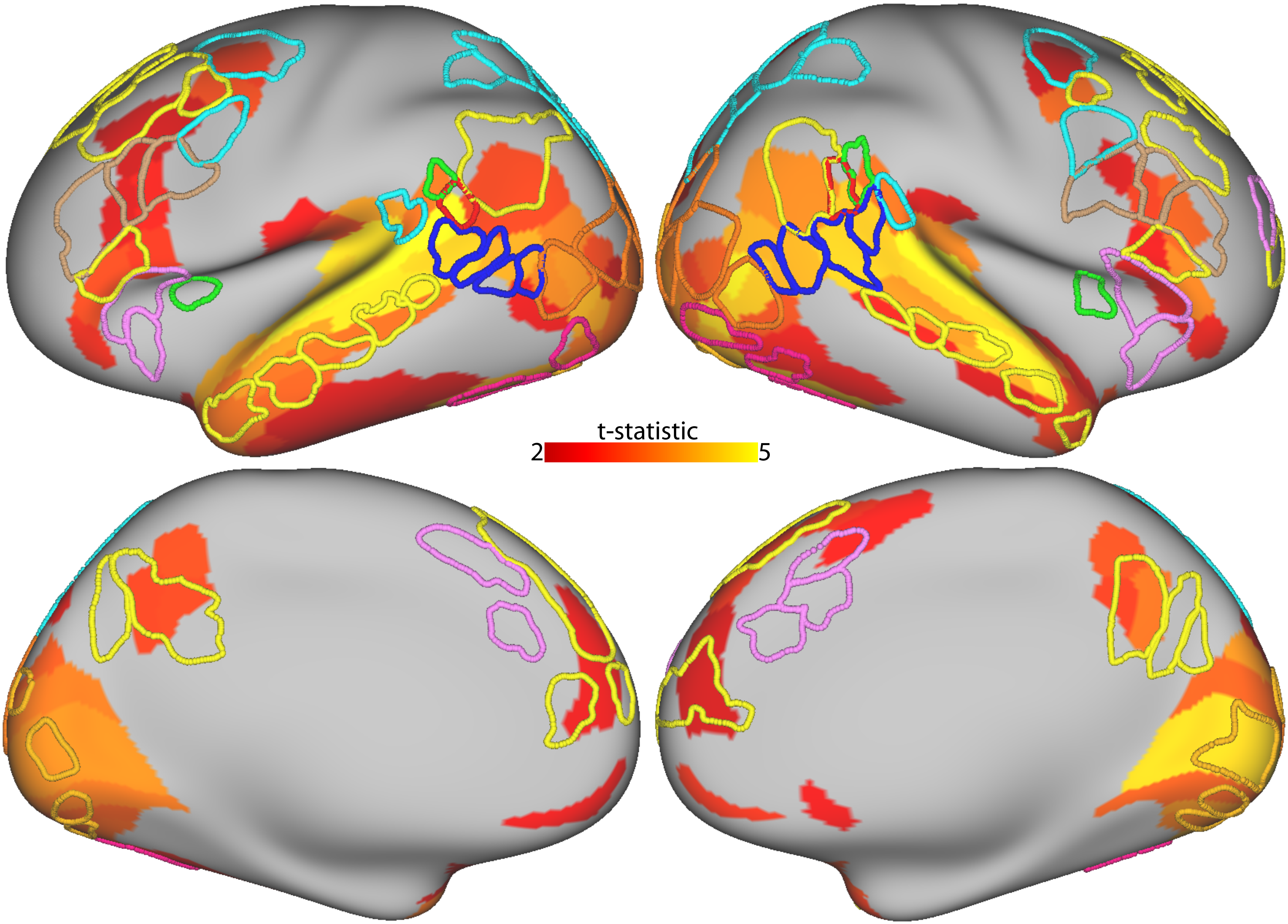

**Supplemental Figure 5**

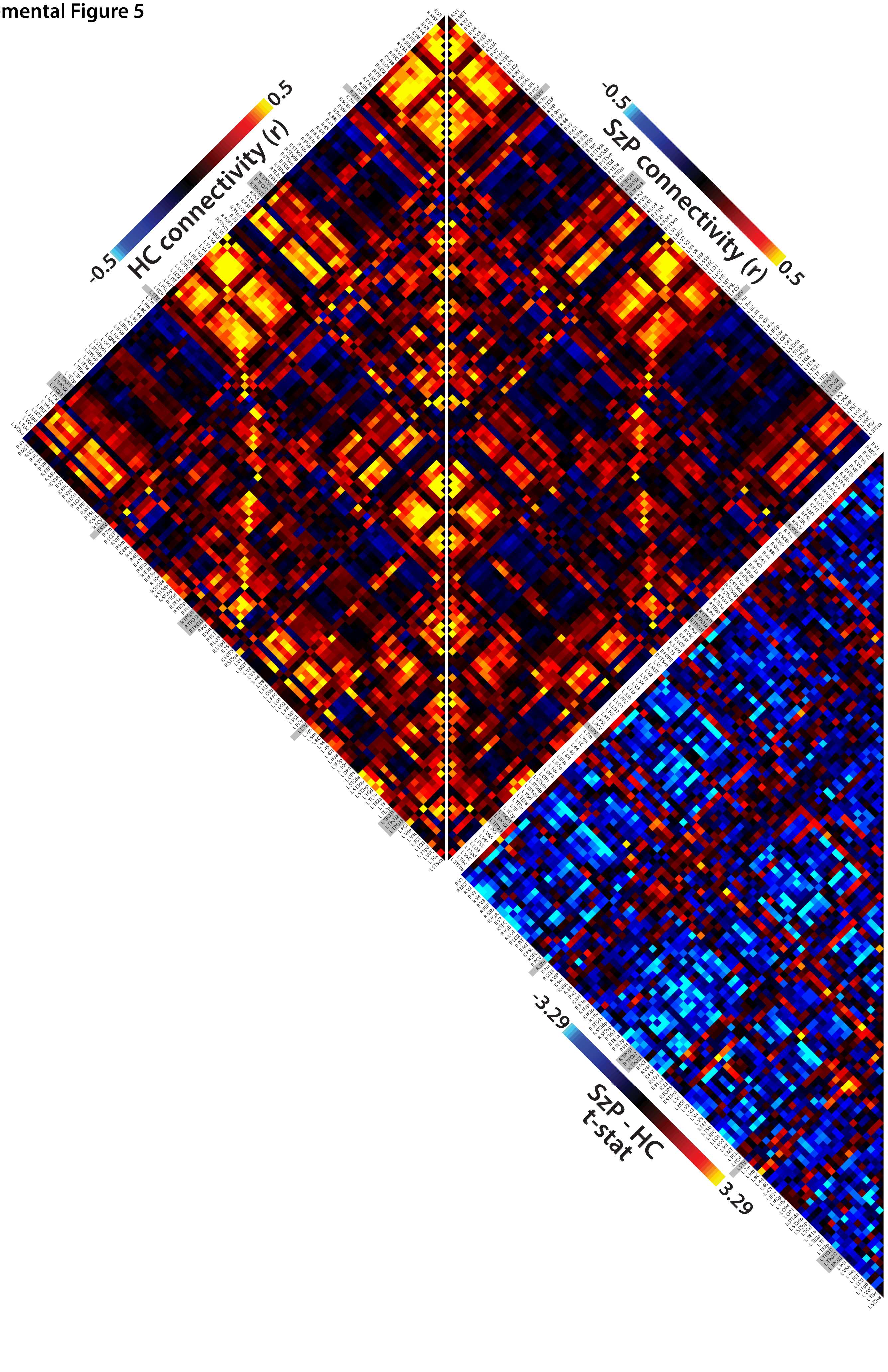

**Supplemental Figure 6**

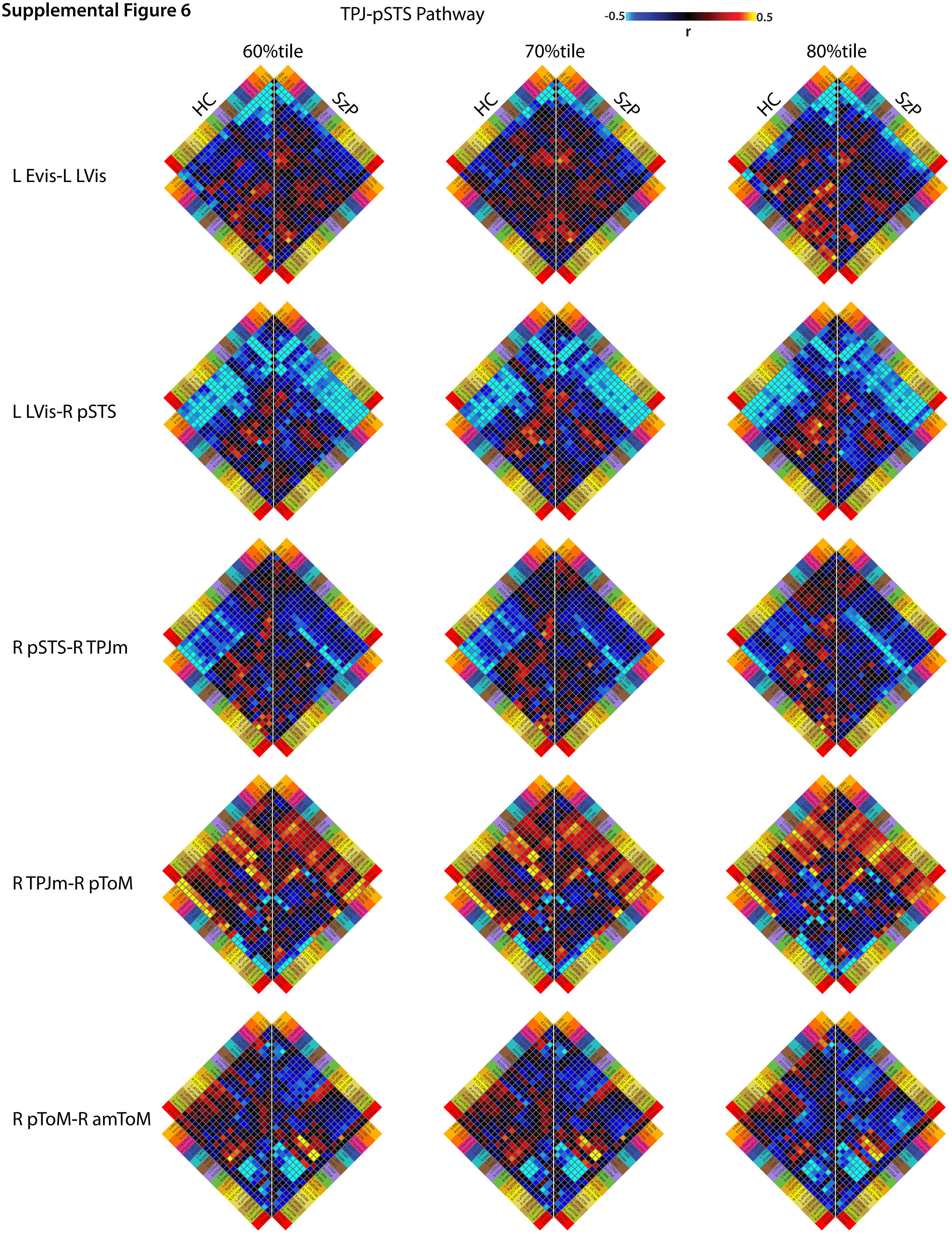

**Supplemental Figure 7**

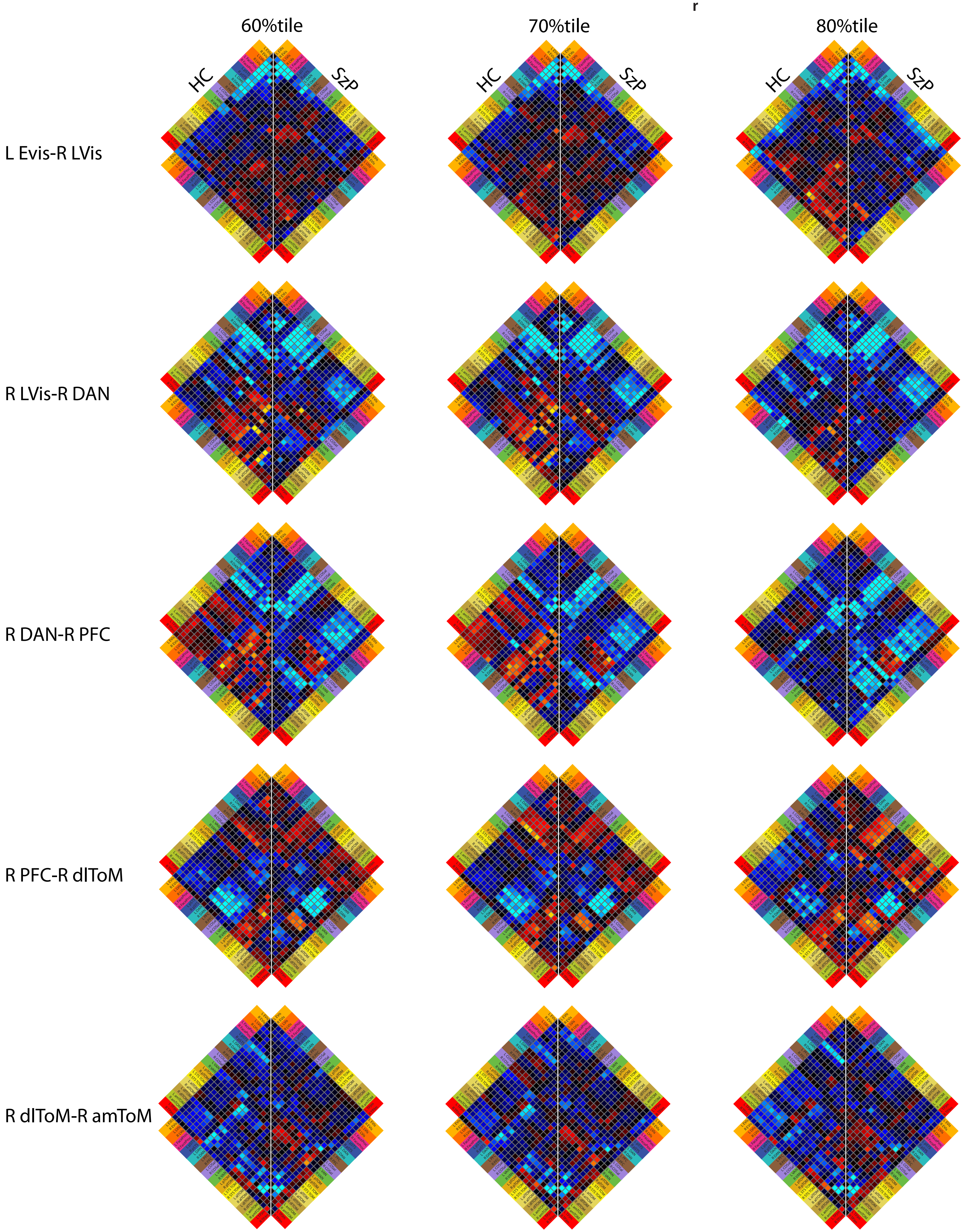

**Supplemental Figure 8**

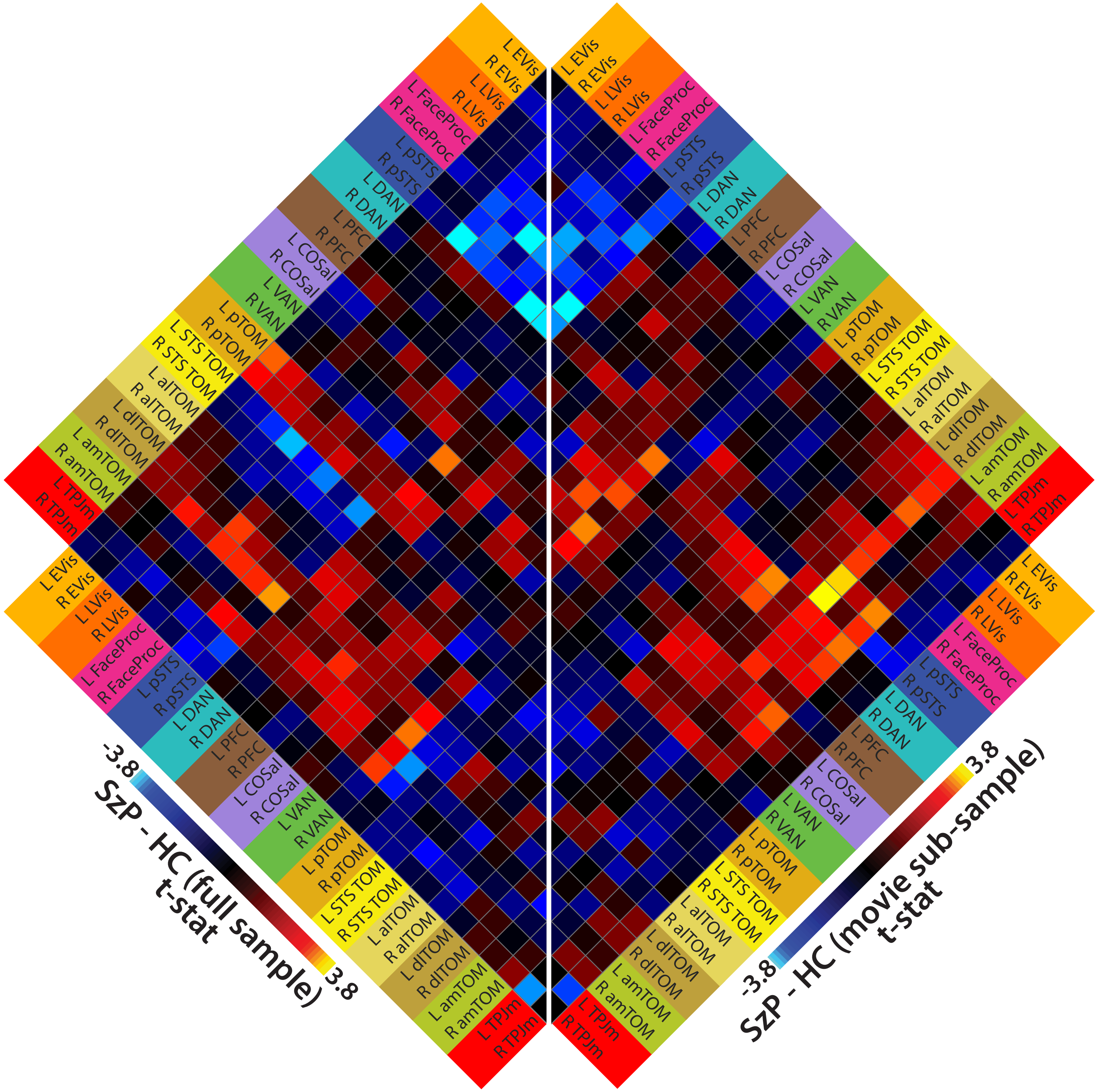

**Supplemental Figure 9**

### A: TPJ-pSTS pathway

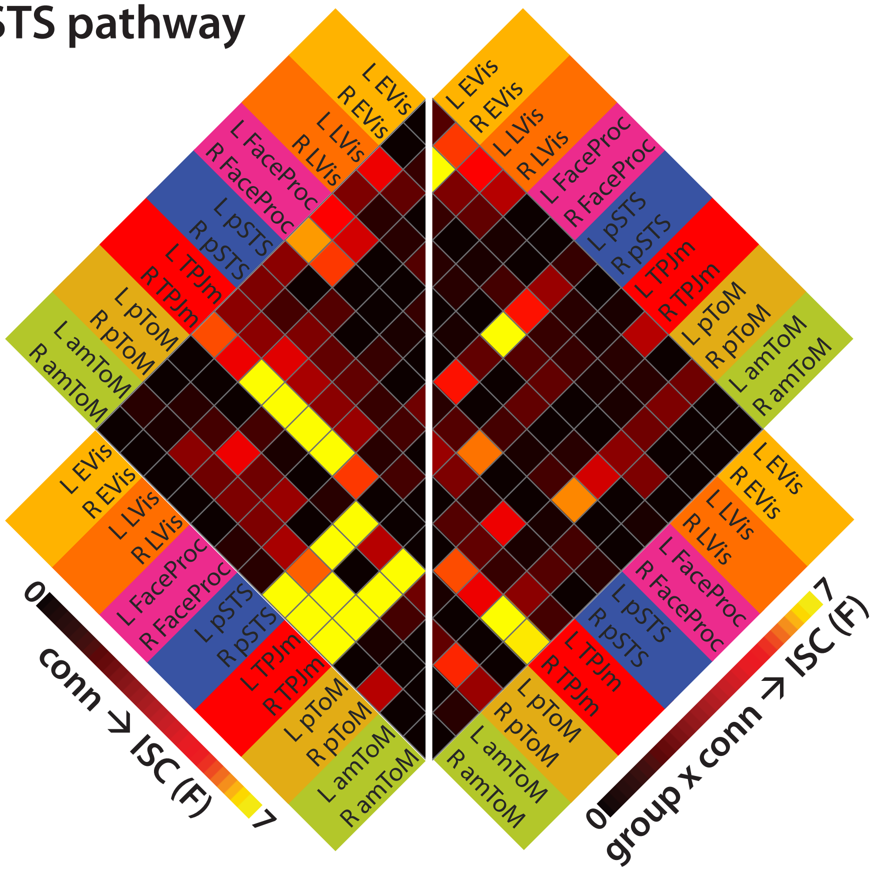


### B: PFC pathway

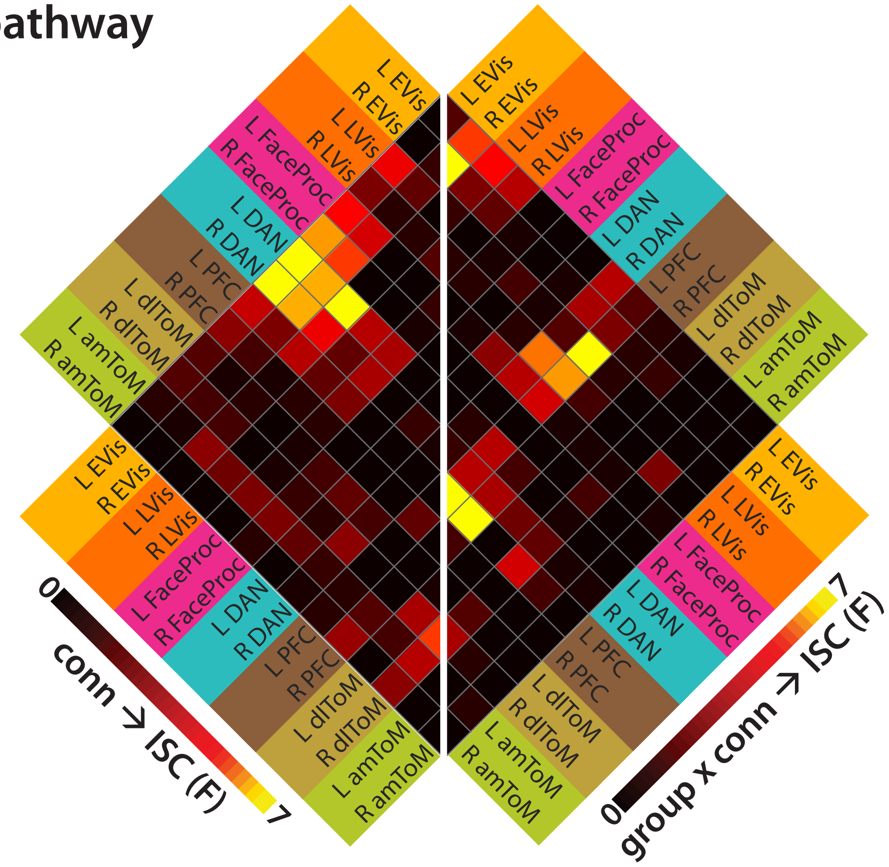
